# Chromatin sequesters pioneer transcription factor Sox2 from exerting force on DNA

**DOI:** 10.1101/2022.02.02.478883

**Authors:** Tuan Nguyen, Sai Li, Jeremy Chang, John Watters, Htet Ng, Adewola Osunsade, Yael David, Shixin Liu

## Abstract

Formation of biomolecular condensates constitutes an emerging mechanism for transcriptional regulation. Recent studies suggest that the co-condensation between transcription factors (TFs) and DNA can generate mechanical forces driving genome rearrangements. However, the reported forces generated by such protein-DNA co-condensation are typically below one piconewton (pN), questioning its physiological significance. Moreover, the force-generating capacity of these condensates in the chromatin context remains unknown. Using single-molecule biophysical techniques, we show that Sox2, a nucleosome-binding pioneer TF, forms co-condensates with DNA, thereby exerting considerable mechanical tension on DNA strands both in *cis* and *trans*. Sox2 can generate forces up to 7 pN—similar in magnitude to other cellular forces. Sox2:DNA condensates are highly stable, withstanding disruptive forces high enough to melt DNA. We find that the disordered domains of Sox2 are required for maximum force generation but not condensate formation per se. Finally, we show that nucleosomes dramatically attenuate the mechanical stress exerted by Sox2 via sequestering it from coalescing on bare DNA. Our findings reveal that TF-mediated DNA condensation can exert significant mechanical stress which can nonetheless be alleviated by the chromatin organization, suggesting a new function of eukaryotic chromatin in protecting the genome from potentially deleterious nuclear forces.

## Introduction

Transcription factors (TFs) bind specific DNA sequences within the genome to regulate the activity of the transcription machinery.^1^ In recent years, a new paradigm for transcriptional control has emerged in which the intrinsically disordered regions (IDRs) of certain nuclear proteins drive the formation of biomolecular condensates and phase-separated sub-compartments.^2–5^ These nuclear compartments, or transcriptional hubs, connect enhancers to promoters, recruit the RNA polymerase and its regulators, and control gene expression in a dynamic fashion.^6,7^ Notably, some TFs have been shown to form co-condensates with DNA,^8–11^ which ensnare an amount of DNA into the condensates and thus exert tension on the outside free DNA.^12^ The force generated by such protein-DNA co-condensation was reported to be in the sub-piconewton (pN) range.^8,9,13^ Whether this mechanical effect driven by TF:DNA co-condensation is strong enough to be relevant in the nuclear milieu remains to be explored. Moreover, the genomic DNA in eukaryotic nuclei is spooled by histone proteins to form nucleosomes and further organized into higher-order chromatin structures.^14^ How chromatin organization impacts TF condensation and its force-generating capacity is still unclear.

In this report, we employed single-molecule imaging and manipulation to directly measure and compare the physical effects of TF:DNA co-condensation on bare and nucleosomal DNA. We chose Sox2 as the model TF, which belongs to the high mobility group (HMG) superfamily, a group of ubiquitous proteins that bind DNA and nucleosomes to induce structural changes in eukaryotic chromatin, initiating cell fate transitions during development and disease.^15^ We observed the real-time formation of Sox2:DNA co-condensates that exert surprisingly high intra-strand and inter-strand mechanical stress. We estimated the maximum condensation force that Sox2 can actively generate to be ~7 pN, an order of magnitude higher than those previously reported for other DNA-binding proteins.^9,13,16^ Remarkably, when nucleosomes were present on DNA, the mechanical effects of Sox2:DNA condensation were drastically reduced. These results suggest that nucleosomes function more than just DNA packaging units, but also as a mechanical barrier to regulate the force generated by protein: DNA co-condensates.

## Results

### Sox2 forms co-condensates with DNA

We used the 48.5 kbp-long bacteriophage λ genomic DNA (λDNA) as a model DNA substrate. λDNA was immobilized on a glass surface via biotin-streptavidin linkage, stained with YOPRO1 fluorescent dye that binds DNA nonspecifically, and imaged with total-internal-reflection fluorescence microscopy (TIRFM) (Fig. 1A). Depending on the separation between the two anchor points, double-tethered λDNA exhibited a distribution of end-to-end distances. Short and slack molecules displayed larger transversal fluctuations than long and taut ones (Fig. 1A). After flowing in 10 nM of Cy5-labeled recombinant full-length human Sox2 (Supplementary Fig. 1), we observed the formation of Sox2 foci on the DNA (Fig. 1B), which endogenously contains numerous cognate Sox2 motifs across its sequence (Supplementary Fig. 2). The brightness of these Sox2 foci varied, but the majority of them clearly harbored more than one molecule based on the observed Sox2 monomer intensities in the same field of view. Notably, Sox2 foci displayed mobility on the DNA as well as multiple fusion and splitting events per recording (Supplementary Fig. 3A), indicating liquid-like properties.^2,17^ Upon Sox2 binding and foci formation, we also observed that the fluorescence signal of the DNA transitioned from a relatively uniform distribution to a few clusters that colocalized with the Sox2 foci (Fig. 1B). This was particularly apparent in the DNA strands with a short tether length.

**Figure 1.**
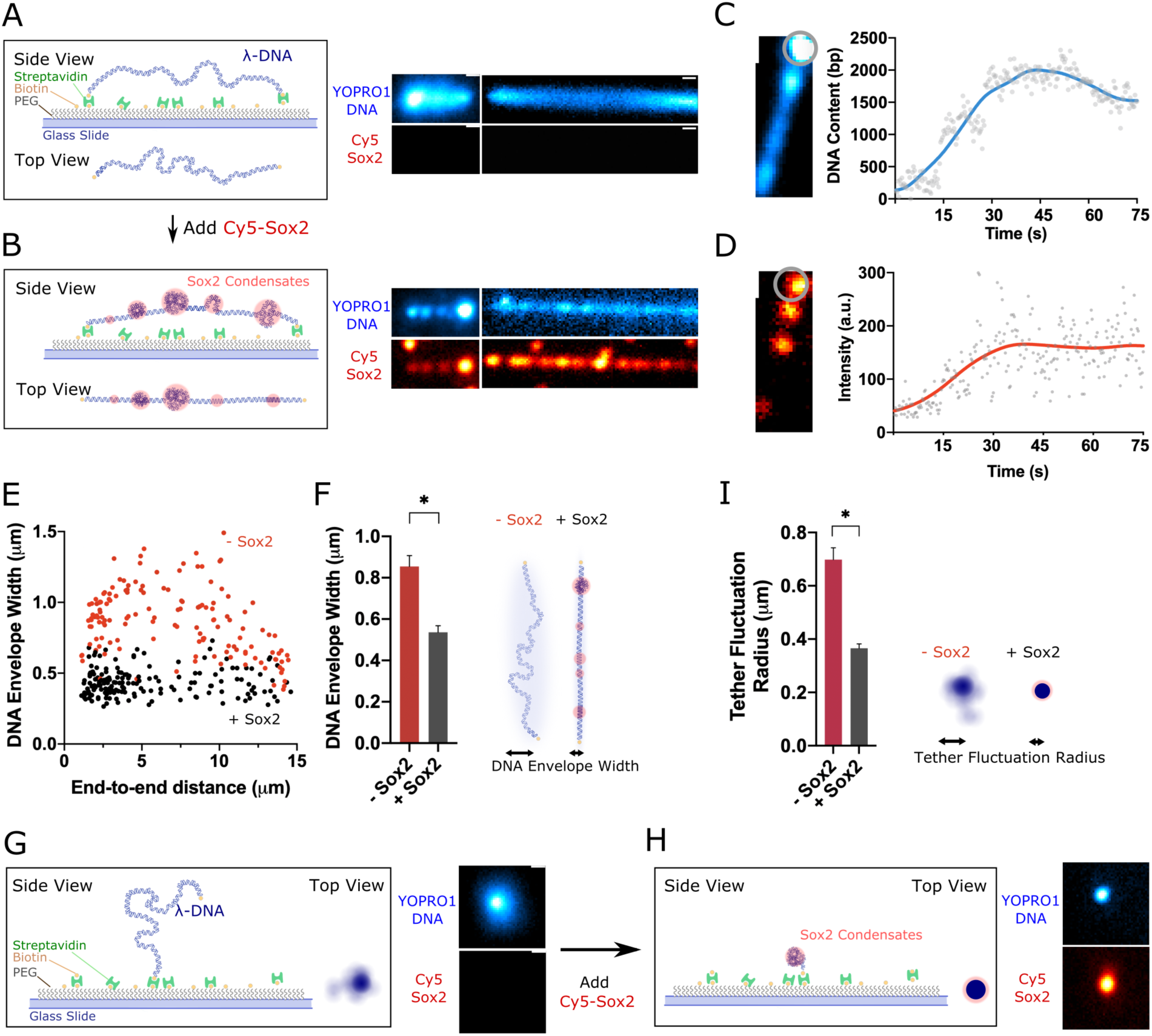
Sox2 forms condensates on DNA. A) (Left) Schematic of a double-tethered λDNA with both ends anchored to a glass surface via biotin-streptavidin linkage. (Right) Two examples of double-tethered λDNA molecules with different end-to-end distances stained with YOPRO1 (20 nM) and imaged by TIRFM. Shown are time-averaged projections over a 75-s period. Scale bar, 0.5 μm. B) Schematic (left) and representative time-averaged projections (right) of double-tethered λDNA (same two molecules as in A) when incubated with 10 nM Cy5-labeled Sox2. Both YOPRO1 and Cy5 channels are shown to illustrate Sox2:DNA co-condensation. C) Real-time tracking of the DNA content (YOPRO1 fluorescence intensity converted to the amount of DNA base pairs) within a condensate (circled region) on a double-tethered λDNA. D) Corresponding changes in the Sox2 intensity within the same circled region as in C. E) DNA envelope width as a function of the end-to-end distance of double-tethered DNA measured in the absence of Sox2 (red) and in the presence of Sox2 (black). *n* = 111, where *n* represents the number of DNA molecules analyzed. F) Bar graph (left) and cartoon (right) showing a reduction in the average DNA envelope width for all the molecules shown in E upon Sox2-mediated co-condensation. G) Schematic (left) and a representative time-averaged projection (right) of single-tethered λDNA stained with YOPRO1 displaying a finite fluctuation radius. H) Schematic (left) and time-averaged projection (right) showing Sox2-mediated condensation of single-tethered λDNA (same molecule as in G). I) Bar graph (left) and cartoon (right) showing a reduction in the average tether fluctuation radius of single-tethered DNA (*n* = 38) upon Sox2-mediated co-condensation.

Once nucleated, the Sox2 foci on DNA were long-lived, and the fluorescence intensities of both Sox2 and DNA at the foci increased with time until reaching a steady state (Fig. 1C and D). Interestingly, we observed a loss of the fluctuating motion in the DNA concurrent with Sox2 foci formation (Fig. 1A and B). Indeed, the average DNA envelope width—a measure for the degree of transversal fluctuations—was significantly reduced in the presence of Sox2 (Fig. 1E and F). Even though the DNA envelope is—as expected—wider for shorter tethers in the absence of Sox2, the addition of Sox2 reduced the envelope width for all double-tethered molecules to the same level (Fig. 1E). These findings can be rationalized by an ability of Sox2 to form co-condensates with DNA. As more DNA being pulled into the condensates, the previously slacked DNA transitioned into a tensed state.

In addition, we observed a fraction of the λDNA that was tethered to the surface at only one end (Fig. 1G), likely because the other biotinylated end did not find a streptavidin to bind during the immobilization step. Without Sox2, these single-tethered DNA molecules displayed random fluctuations characterized by a finite radius (Fig. 1G and I). The addition of Sox2 again visibly suppressed such fluctuations (Supplementary Video 1), most likely due to co-condensation, resulting in a significantly decreased average fluctuation radius (Fig. 1H and I). Altogether, these results demonstrate that Sox2 and DNA form co-condensates wherein proteins and DNA accumulate, reducing the amount of free DNA outside the condensates.

### Sox2:DNA co-condensation exerts high tension on DNA

The loss of fluctuations in both single- and double-tethered DNA suggests that Sox2-induced condensation generates mechanical tension within the DNA. In accordance with this notion but nonetheless unexpectedly, we observed that a significant population of double-tethered DNA underwent sudden breakage after losing slacks (Fig. 2A and B, Supplementary Video 2). The breakage was accompanied by a rapid collapse of the fluorescence signals from the DNA and Sox2 into each tethered end (Supplementary Video 2). Importantly, these breakage events occurred over a time window that coincided with the formation of Sox2:DNA co-condensates (Supplementary Fig. 3B). In contrast, minimal DNA breakage was observed in the absence of Sox2 (Fig. 2B). The fraction of broken DNA tethers was not significantly affected by the concentration of YOPRO1 or the duration of laser exposure in the experiments (Supplementary Fig. 4A and B), arguing against the possibilities of dye- or laser-induced DNA structural instability being the main source of DNA breakage. Moreover, under the same imaging condition, we detected virtually no breakage or dissociation of single-tethered λDNA—where the tension can be released from the free end—after the addition of Sox2 (Supplementary Fig. 4C). These observations suggest that Sox2-mediated DNA condensation generates considerable tension within the DNA when both ends are anchored.

**Figure 2.**
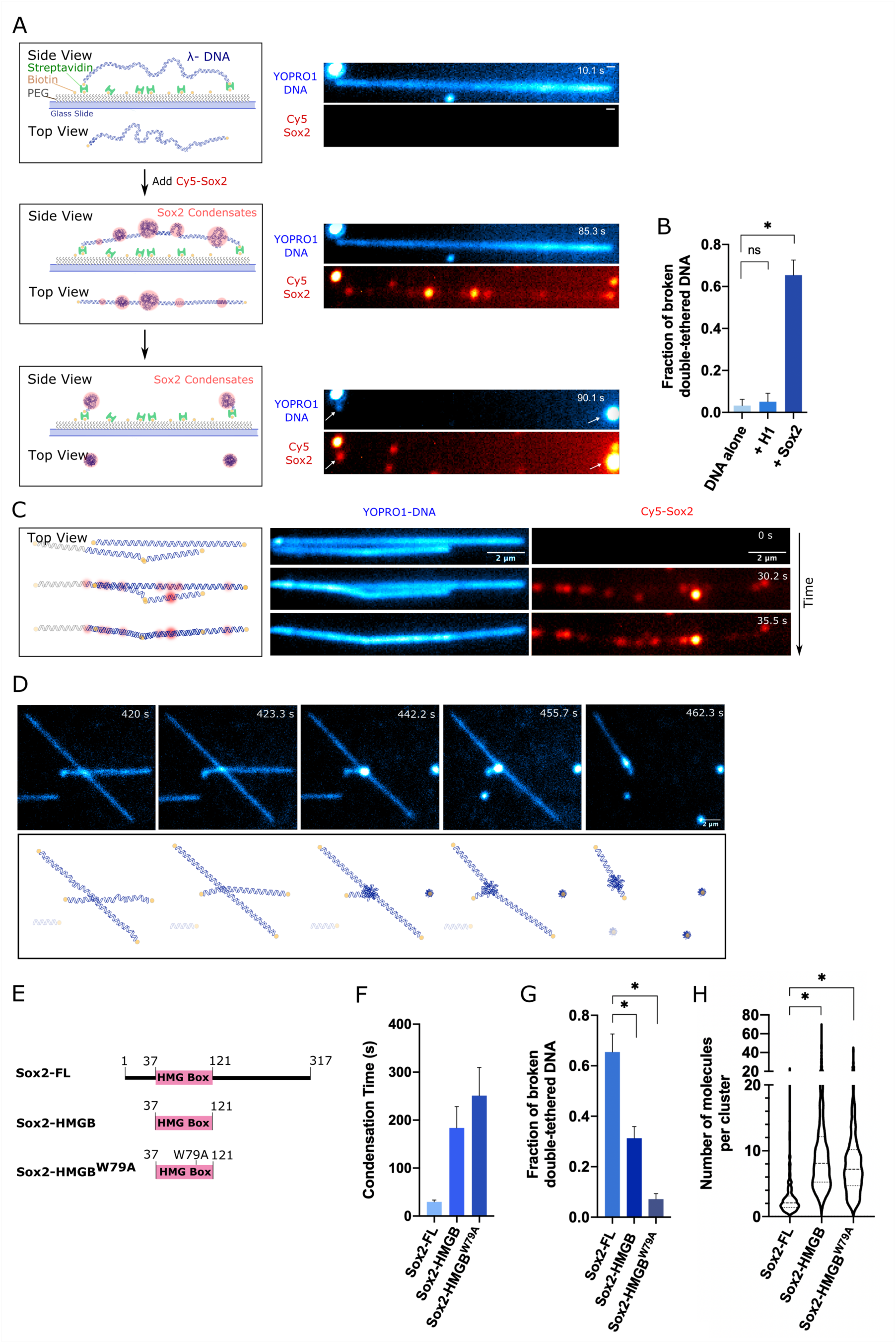
Sox2:DNA co-condensation exerts intra-strand and inter-strand force. A) Schematic (left) and time-lapse snapshots (right) of Sox2 condensate formation on a double-tethered λDNA and the subsequent breakage event upon which both DNA and Sox2 signals collapsed to the two tethered ends (white arrows). Scale bar, 0.5 μm. B) Fraction of double-tethered λDNA molecules that broke after 15 min without any protein (*n* = 251), with H1 (*n* = 150), and with Sox2 (*n* = 379). Data are averaged from at least three fields of view. Error bars denote 95% confidence intervals. C) Schematic (left) and time-lapse snapshots (right) showing multiple adjacent DNA strands being joined by Sox2-mediated condensation. C) Time-lapse snapshots (top) and cartoon illustrations (bottom) of a sequence of DNA breaking and joining events occurring among multiple λDNA strands in the presence of Sox2. E) Schematic of different Sox2 constructs used in this study. F) Average condensation time from at least three fields of view for each Sox2 construct. Error bar denotes standard deviation. G) Fraction of double-tethered λDNA molecules that broke after 15 min of incubation with Sox2-FL (*n* = 379), Sox2-HMGB (*n* = 357), or Sox2-HMGB^W79A^ (*n* = 297). H) Violin plot showing the distribution of the number of Sox2 molecules within each cluster for Sox2-FL (*n* = 381), Sox2-HMGB (*n* = 937), or Sox2-HMGB^W79A^ (*n* = 546), where *n* represents the number of clusters analyzed.

To examine whether other DNA-binding proteins can exert the same level of tension on DNA, we repeated the above TIRFM assay with another abundant nuclear protein, the human linker histone H1.4 (referred to as H1 hereafter). We found that H1 also formed co-condensates with DNA,^10^ which reduced the envelope width for double-tethered DNA and fluctuation radius for single-tethered DNA (Supplementary Fig. 5). However, H1-mediated DNA condensation resulted in much fewer DNA breakage events compared to Sox2-mediated condensation (Fig. 2B), indicating that H1 generates a lower force on DNA, consistent with previous results.^8^

We then sought to examine whether Sox2:DNA co-condensation can generate inter-strand tension by observing neighboring λDNA strands immobilized in close proximity of each other. In the absence of Sox2, the DNA molecules fluctuated independently. Upon the addition of Sox2, these strands lost slack and joined one another through the fusion of Sox2 foci (Fig. 2C and Supplementary Video 3). In some cases, we observed successive severing and joining of DNA located nearby (Fig. 2D and Supplementary Video 4). Together, these results suggest that Sox2 condensates exert force on DNA both within the same strand and between multiple strands.

### IDRs of Sox2 are dispensable for condensate formation but required for force exertion

To gain insights into the structural domains driving the capacity of Sox2 to form co-condensates with DNA, we generated and fluorescently labeled Sox2 truncations (Fig. 2E and Supplementary Fig. 1). Sox2 contains N- and C-terminal IDRs flanking the globular DNA-binding HMGB domain.^18^ We first examined a Sox2 construct that contains only its HMGB domain without the IDRs (Sox2-HMGB). Unexpectedly, similar to the full-length Sox2 (Sox2-FL), Sox2-HMGB also formed foci on λDNA strands—both doubly and singly tethered—along with a concomitant loss of DNA fluctuations (Supplementary Fig. 6). This observation indicates that the IDRs of Sox2 are not required for its co-condensation with DNA. However, Sox2-HMGB took ~5 times longer to form the same amount of DNA co-condensates (measured through the loss of fluctuations) compared to Sox2-FL (Fig. 2F). Sox2-HMGB-mediated DNA condensation also resulted in significantly fewer DNA breakage events (Fig. 2G). We next introduced a single-residue mutation, W79A, to Sox2-HMGB (Sox2-HMGB^W79A^). Consistent with a previous report,^19^ Sox2-HMGB^W79A^ displayed diminished DNA binding activity (Supplementary Fig. 7). However, it still retained co-condensation activity with DNA (Supplementary Fig. 6), albeit with much slower condensation kinetics compared to Sox2-FL (Fig. 2F). This mutation further attenuated the condensation-dependent mechanical tension exerted on DNA (quantified by the frequency of double-tethered DNA breakage) compared to Sox2-FL and Sox2-HMGB (Fig. 2G). Notably, the mechanical effect of Sox2:DNA co-condensates is not directly correlated with their size, as both Sox2-HMGB and Sox2-HMGB^W79A^ foci on average contained a higher number of Sox2 molecules compared to Sox2-FL ones (Fig. 2H). Together, these results demonstrate that HMGB alone can mediate Sox2:DNA co-condensation, but the strong mechanical pressure on DNA is largely driven by the IDRs of Sox2.

### Quantification of the forces generated by Sox2:DNA co-condensation

Next, we sought to quantitatively measure the force exerted by Sox2:DNA co-condensates on the DNA strand. Using optical tweezers combined with scanning confocal microscopy, we tethered a single λDNA molecule between two optically trapped beads, moved the tether in a relaxed form (i.e. zero applied force) to a channel containing Cy3-labeled Sox2, and monitored the force on DNA as a function of time in a passive mode (i.e. keeping the trap positions fixed), while simultaneously observing condensate formation (Fig. 3A). We observed that Sox2 foci appear on the DNA tether within a few seconds after moving to the Sox2 channel (Supplementary Fig. 8) and a concurrent increase in force that plateaued at ~7 pN (Fig. 3B).

**Figure 3.**
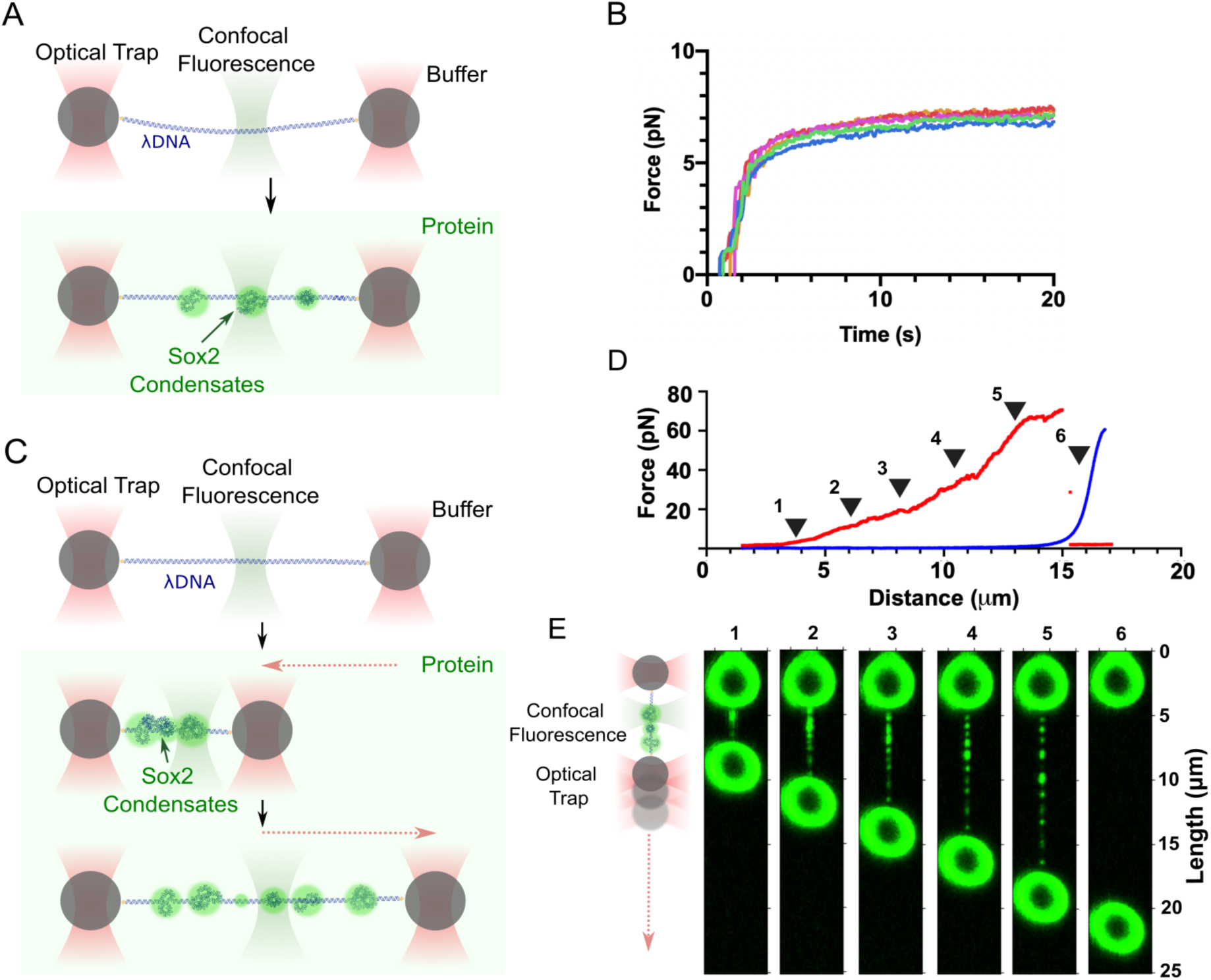
Sox2:DNA co-condensates are mechanically stable. A) Schematic of the optical tweezers assay that measures forces generated by Sox2-mediated DNA condensation. The trap positions were held fixed in this experiment. B) Representative force measurements as a function of time for the assay depicted in A. Each colored line represents an individual λDNA tether on which Sox2 condensates formed. C) Schematic of the optical tweezers assay in which Sox2 condensates were first allowed to form on λDNA under a 0.5-pN force clamp—causing tether shortening—and then subjected to mechanical pulling by gradually separating the two traps apart. D) A representative force-distance curve from pulling a λDNA tether harboring Sox2 condensates (red) in comparison to a representative curve from pulling a bare λDNA (blue). E) 2D fluorescence scan of the same tether as in D (red) at different time points showing Sox2 condensates persisting under an increasing force up to 60 pN (time points 1-5) until tether breakage (time point 6).

We then conducted force-clamp experiments in which Sox2 was incubated with tethered DNA maintained at a specified force value (Supplementary Fig. 9A). We observed that, at a force clamp of 0.5 pN, Sox2 and DNA underwent continued condensation, which reduced the length of free DNA and brought the two beads closer to each other (Supplementary Fig. 9B). In contrast, a 10-pN force clamp largely abrogated the condensation process (Supplementary Fig. 9C). These results are in agreement with the above passive-mode results reporting a 7-pN maximum force that Sox2:DNA co-condensates can generate (Fig. 3B).

Next, we asked how much force is required to dissolve Sox2:DNA co-condensates. To address this question, we first formed Sox2 foci on a DNA tether under a 0.5-pN force clamp and then gradually pulled the two beads apart, thereby increasing the force applied to the tether (Fig. 3C). From the resultant force-extension curve, we found that the extension of a Sox2-bound tether was much shorter than that of a bare DNA tether, indicating significant DNA accumulation inside the condensates (Fig. 3D). Some rupture events were observed with increased force, suggesting condensate dissolution (Fig. 3D). Nonetheless, a significant fraction of condensates remained intact even when the force reached the DNA overstretching regime (~65 pN), as reflected by the shorter extension at high forces compared to bare DNA (Fig. 3D). Concomitant fluorescence scanning confirmed persistence of condensates during pulling until tether breakage (Fig. 3E). These results demonstrate that Sox2:DNA co-condensates are stable and resistant to high disruptive forces.

### Nucleosomes attenuate Sox2-driven mechanical effects on DNA

The high forces that Sox2 condensates exert on DNA raises the question of how this process is regulated in cells to avoid unwanted genome instability. We surmised that the eukaryotic chromatin may serve as a barrier against such mechanical stress, particularly given that Sox2 is a nucleosome-binding pioneer TF.^20^ To test this hypothesis, we loaded histone octamers containing Cy3-labeled H2B onto surface-immobilized λDNA in the TIRFM setup (Fig. 4A), and incubated the nucleosomal DNA with Cy5-Sox2 (Fig. 4B). We observed Sox2 foci nucleate around nucleosome locations (Fig. 4C and D). Sox2 foci preferentially colocalized with nucleosomes over non-nucleosomal DNA sites (Fig. 4E and F). The majority of Sox2:nucleosome foci contained multiple Sox2 molecules based on the Cy5 fluorescence intensity, similar to Sox2 foci on bare DNA (Supplementary Fig. 10A).

**Figure 4.**
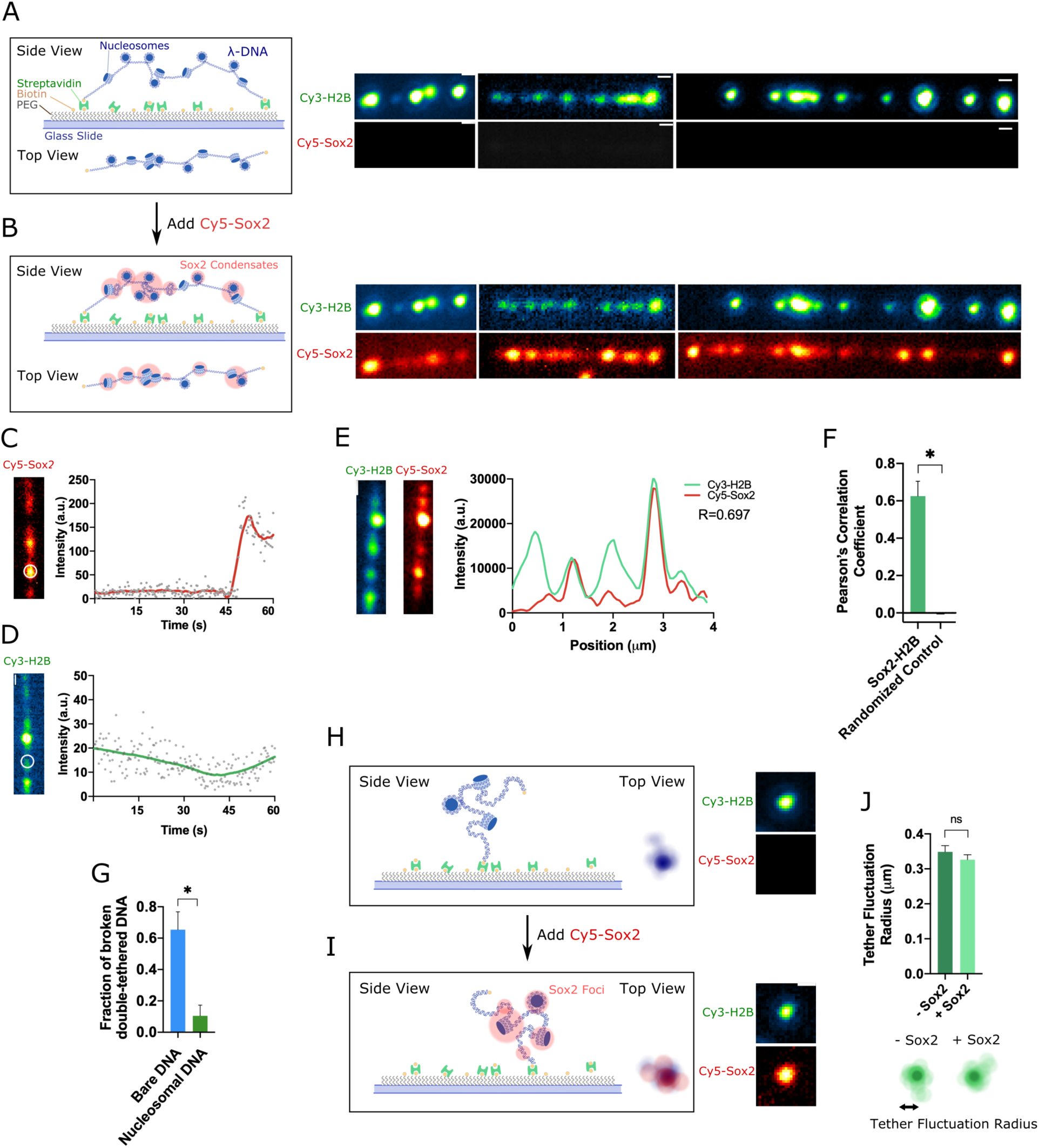
Nucleosomes colocalize with Sox2 condensates and attenuate their mechanical effects on DNA. A) (Left) Schematic of double-tethered λDNA loaded with nucleosomes. (Right) Time-averaged projections of three double-tethered nucleosomal DNA molecules containing Cy3-H2B with different end-to-end distances. Scale bar, 0.5 μm. B) Schematic (left) and representative time-averaged projections (right) of double-tethered nucleosomal λDNA containing Cy3-H2B (same three molecules as in A) when incubated with 10 nM Cy5-labeled Sox2. C) Real-time tracking of Cy5-Sox2 intensities at a nucleosome position (circled region) on a double-tethered nucleosomal λDNA. D) Corresponding Cy3-H2B intensities within the same circled region as in C. E) (Left) Snapshot of a double-tethered DNA harboring Cy3-H2B nucleosomes and Cy5-Sox2 condensates. (Right) Intensity profiles of Cy3-H2B (green) and Cy5-Sox2 (red) along the length of the same DNA molecule. *R* value represents the Pearson’s correlation coefficient. F) Average Pearson’s correlation coefficients for all aligned Cy3-H2B and Cy5-Sox2 intensity profiles and for Costes’ randomized control (*n* = 158). G) Fraction of double-tethered bare DNA (*n* = 379) versus nucleosomal DNA molecules (*n* = 303) that broke after 15 min of incubation with 10 nM Sox2. Data are averaged from at least three fields of view. H) Schematic (left) and a representative time-averaged projection (right) of a single-tethered nucleosomal λDNA (visualized by Cy3-H2B fluorescence). I) Schematic (left) and time-averaged projection (right) of the same single-tethered nucleosomal DNA molecule as in H when incubated with 10 nM Cy5-Sox2. J) Average tether fluctuation radius of single-tethered nucleosomal λDNA in the absence and presence of Sox2 (*n* = 62).

Strikingly, we detected drastically fewer DNA breakage events upon the formation of Sox2 foci on nucleosomal DNA compared to those on bare DNA (Fig. 4G). In a few examples in which nucleosomal DNA breakage was observed, it appeared that the tether broke at the end, and the full contour of the nucleosomal λDNA was maintained and underwent rigid-body fluctuations (Supplementary Fig. 10B and Supplementary Video 5). This is in contrast to the breakage events observed on bare DNA where the tether broke in the middle and the Sox2 and DNA signals abruptly collapsed into opposite ends (Fig. 2A and Supplementary Video 2).

We interpreted these results as Sox2 molecules being largely sequestered at the nucleosomal sites and thus failing to reel in free DNA and generate large forces. We further analyzed the single-tethered nucleosomal λDNA that displayed randomly fluctuating motions. The addition of Sox2 did not significantly suppress the fluctuations of nucleosomal DNA (Fig. 4H-J and Supplementary Video 6), in contrast to single-tethered bare DNA results (Fig. 1I and Supplementary Video 1). Together, these results suggest that nucleosomes attenuate the mechanical stress exerted by Sox2 on DNA through colocalization with Sox2 (Fig. 5).

**Figure 5.**
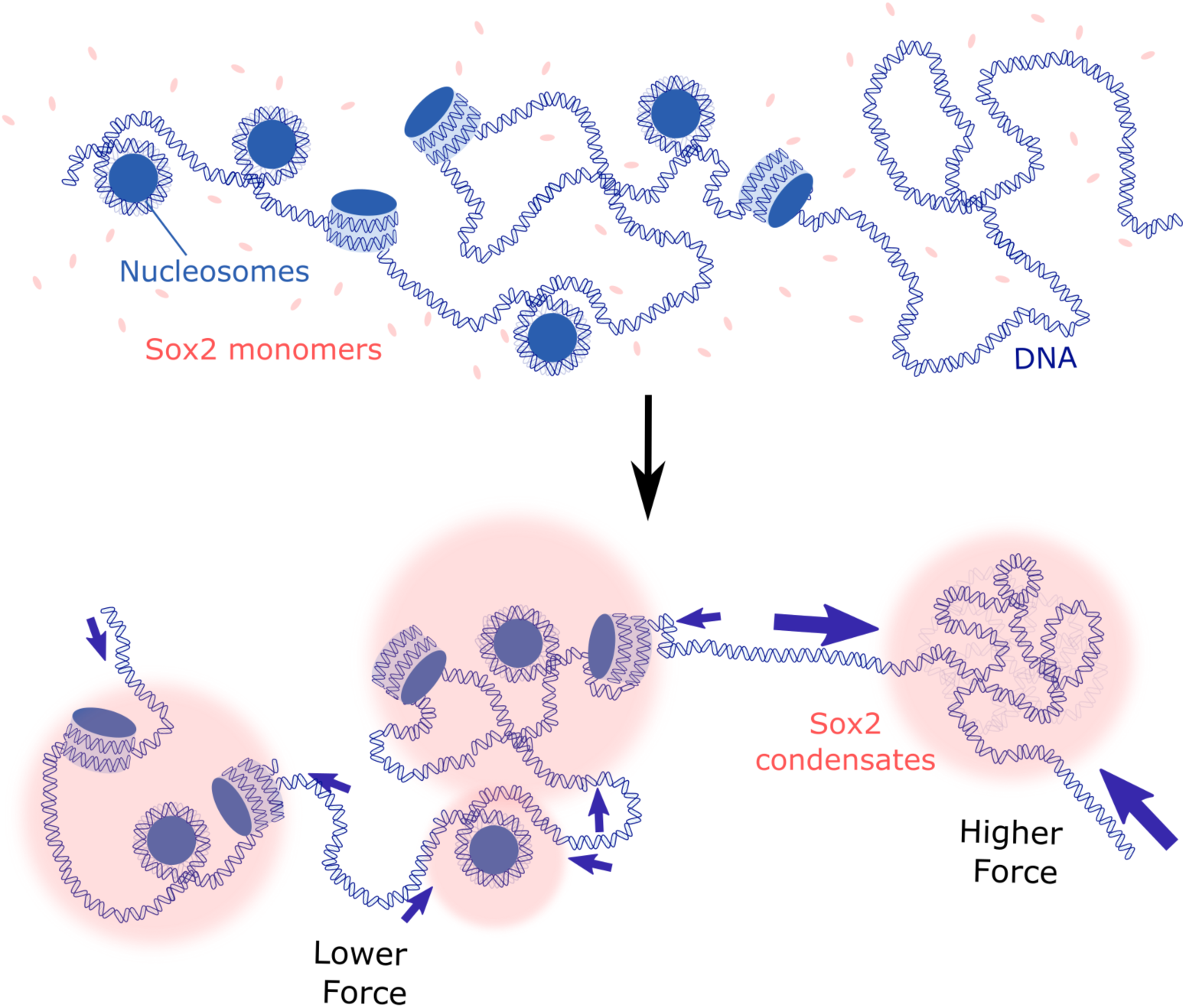
Model depicting the distinct mechanical effects of Sox2-mediated condensation on nucleosomal DNA versus nucleosome-free DNA.

## Discussion

Prior to this study, the forces generated by co-condensation between DNA and proteins—such as FoxA1 and PARP1—were estimated to be on the order of sub-pN, placing them among the weakest nuclear forces alongside those generated by loop-extruding SMC complexes such as condensin and cohesin.^9,13,21^ Here we show that Sox2, an abundant TF central to pluripotency and embryogenesis, can actively generate condensation forces up to 7 pN, one order of magnitude higher than previously reported values. It is worth noting that Klf4, another pluripotency TF, can also form condensates on DNA against a relatively high force (~8 pN) according to a recent report.^22^ Once formed, the Sox2:DNA co-condensates are extremely stable, resisting against pulling forces sufficient to overstretch B-form DNA. In comparison, a fraction of the condensates formed by DNA and Heterochromatin Protein 1α can resist disruptive forces of up to 40 pN, while others rupture at lower forces between 5 and 20 pN.^23^ These findings are of significance because they show that forces exerted by certain protein:DNA co-condensates are comparable to other cellular forces such as those generated by many molecular motors.^24^ Therefore, TF-mediated DNA condensation can potentially impact other genomic processes by maintaining a mechanically stable sub-compartment. We speculate that the range of forces displayed by different protein:DNA co-condensates represent factor-specific modes of gene regulation that can be further tuned *in vivo* to achieve spatiotemporal control.

The critical force below which a protein-rich condensate is able to pull DNA inside likely depends on the physicochemical properties of the condensate, such as its surface tension,^25^ which in turn are determined by the characteristics of the TF including its charge distribution and intrinsic disorder. In this work, we show that the ability of Sox2 to generate high forces through co-condensation with DNA critically relies on its IDRs; on the other hand, the DNA-binding HMGB domain of Sox2 alone is sufficient for condensation with DNA. This is reminiscent of recent findings with Klf4 and SMN proteins,^26,27^ suggesting that condensate formation and force generation are two separable activities dependent on distinct domains of a given TF.

Eukaryotic chromatin is known to profoundly alter the mechanics of DNA by changing its persistence length and torsional stiffness.^28,29^ Our work presented here adds to the mechanical regulatory roles of chromatin by showing that nucleosomes sequester TFs such as Sox2 from exerting high mechanical tension on DNA by forming co-condensates. Considering that Sox2 has the inherent ability to bind nucleosomes, it is conceivable that such interactions serve as nucleation events that eventually recruit most of the available Sox2 molecules to nucleosomal sites. These Sox2:nucleosome condensates differ from the Sox2:DNA ones in their physicochemical properties, thereby resembling a mechanical “sink” that buffers the stress within genomic DNA (Fig. 5). Whether this effect is limited to pioneer transcription factors is an interesting question to answer in the future.^30^

In sum, our study highlights the mechanical impact of TF:DNA co-condensation on the genome and the role of chromatin in regulating such mechanical stress. It can be envisioned that the nucleosome landscape—shaped by many factors and altered in developmental and disease states—is directly related to the force field in the nucleus. Further studies are warranted to elucidate this relationship, which will improve our understanding of how chromatin mechanics influence genome architecture and gene expression.^31^

## Acknowledgements

We thank members of the Liu Laboratory for discussions, R. Shih (Liu Laboratory) and the O’Donnell Laboratory (Rockefeller University) for protein reagents, and Rockefeller University Bio-Imaging Resource Center for help with image analysis. T.N. and J.C. are supported by a Medical Scientist Training Program grant from the National Institute of General Medical Sciences of the National Institutes of Health under award number T32GM007739 to the Weill Cornell/Rockefeller/Sloan Kettering Tri-Institutional MD/PhD Program. Y.D. is supported by NIH grant R35GM138386 and CCSG core grant P30CA008748. S.Liu is supported by the Robertson Foundation, the Pershing Square Sohn Cancer Research Alliance, and an NIH Director’s New Innovator Award (DP2HG010510).

## Author Contributions

S.Liu and T.N. conceived the project and designed the experiments. S.Li cloned the Sox2 constructs and prepared the histone octamers. T.N. purified and labeled Sox2 proteins, performed TIRFM experiments, as well as bulk assays with help from H.N.. J.C. performed the optical tweezers experiments. J.W. wrote scripts for data analysis. A.O. and Y.D. provided the linker histones. T.N. and S.Liu wrote the manuscript with input from all authors.

## Methods

### Protein purification and labeling

#### Sox2

Human Sox2 proteins were expressed and purified as previously described.^32^ In brief, Sox2-FL and Sox2-HMGB constructs were cloned into the pET28B plasmid, expressed in Rosetta (DE3) plyS cells (Novagen #70956-3) in LB media at 37°C until reaching an OD_600_of ~0.6, and induced with 0.5 mM IPTG at 30 °C for 2 h. Cells were harvested, lysed, and purified using Ni-NTA affinity column under denaturing condition. Eluted Sox2 was refolded by changing to a zero-urea buffer using a desalting column (GE healthcare #17-1408-01). Further purification was performed by gel filtration on a Superdex 200 10/300 GL column (GE Healthcare). Fluorescence labeling was performed as previously described.^32^ In brief, Cy5 or Cy3 maleimide (GE healthcare) was mixed with Sox2 at a molar ratio of ~2:1. For Sox2-FL, the dye was conjugated to the only native cysteine C265. For Sox2-HMGB, a K42C mutation was introduced by site-specific mutagenesis. Free dye was removed by gel filtration on a Superdex 200 10/300 GL column.

#### Histone octamer

Recombinant histone octamers from *Xenopus laevis* were purified and labeled as previously described^32^. In brief, each of the four core histones was individually expressed in BL21 (DE3) cells, extracted from inclusion bodies, and purified under denaturing conditions using Q and SP ion exchange columns (GE Healthcare). Octamers were refolded by dialysis and purified by gel filtration on a Superdex 200 10/300 GL column. To label the octamer, single-cysteine construct H2B T49C was generated by site-directed mutagenesis and incubated with Cy3 maleimide at 1:5 molar ratio.

#### Linker histone H1

His-Sumo-H1.4^A4C^-GyrA-His was expressed and purified as described previously^33^ with minor adjustments. Briefly, the construct was expressed in Rosetta DE3 cells overnight at 16^°^ C. Cells were lysed and lysate incubated with Ni-NTA beads (Bio-Rad). 1 mM DTT was added to the eluent, and it was incubated with Ulp-1 (1:100 v/v) for 1 h at room temperature. Following this, 500 mM BME was added. The mixture was run on a Hi-Trap SP column, and fractions containing full-length H1.4^A4C^ were pooled and injected on a semi-preparative HPLC C18 column. Pure fractions of H1.4^A4C^ were pooled and lyophilized. Lyophilized H1.4^A4C^ was resuspended in H1 labeling buffer (6 M Guanidine, 20 mM Tris pH 7.5, 0.2 mM TCEP). It was mixed with 3 molar equivalents of Cy3 maleimide for 1 h at room temperature, followed by quenching with 1 mM BME. This was injected on a semi-preparative HPLC C18 column. Pure fractions of Cy3-H1.4 were pooled and lyophilized. Cy3-H1.4 was resuspended in H1 buffer (20 mM Tris pH 7.5, 200 mM NaCl) before use.

### Single-molecule TIRFM experiments

Single-molecule imaging was conducted on a total-internal-reflection fluorescence microscope (Olympus IX83 cellTIRF). PEG slides were prepared as previously described.^32^ The assembled flow chamber was infused with 20 μL of 0.2 mg/mL streptavidin (Thermo Fisher Scientific),incubated for 5 min, and washed with 250 μL of T150 buffer (50 mM Tris pH 7.5, 150 mM NaCl, 0.0075% Tween). Biotinylated λDNA (LUMICKS) was immobilized by slowly injecting a diluted 10-20 pM solution at a volume of 40-80 μL over the course of 2 min, which was achieved using a pipette or a pump. 250 μL of T150 buffer was flowed into the injection chamber to wash away molecules that were not immobilized. 100 μL of a solution containing DNA staining dye and imaging buffer components [T150 buffer, 20 nM YOPRO1, oxygen scavenging system (4% w/v glucose, 1.5 mg/mL glucose oxidase, 0.072 mg/mL catalase, 2 mM Trolox)] was flowed in to visualize immobilized λDNA. In the nucleosome experiments, we adopted a previously described protocol with minor modifications.^34^ In brief, *in situ* nucleosome formation was achieved via flowing in Cy3-labeled histone octamer with Nap1 (provided by O’Donnell lab and R. Shih) into the chamber followed by a 5-min incubation and a washing step with T150. After that, a solution cocktail containing 10 nM of Cy5-labeled Sox2 and the above imaging components (i.e. T150, YOPRO1, and oxygen scavenging system) were prepared, 50 μL of which was flowed into the microfluidic chamber, and movies/images were recorded. H1 imaging was similarly performed with 150 pM of Cy3-H1 being flowed in with 30 nM of TOTO3 intercalating dye. Movies were recorded at room temperature with a frame rate of 300 ms. 488-nm, 532-nm, and 640-nm lasers were used to illuminate YOPRO1, Cy3, and Cy5/TOTO3 dyes, respectively. Movies were subsequently displayed and analyzed using plugins in ImageJ/FIJI.

### TIRFM data analysis

#### Analysis of DNA envelope width and fluctuation radius

We followed a general pipeline as previously described.^13^ In brief, we generated time-averaged projections of DNA images in conditions with/without proteins. Transverse line profile of the DNA intensity was generated by drawing a line perpendicular to the middle of the DNA, which gives the maximum DNA width. Background was subtracted off these profiles, and a Gaussian curve was fitted to each line profile. The DNA envelope width and fluctuation radius were defined as two times the standard deviation of the fitted Gaussian curve.

#### Estimation of DNA content and Sox2 counts in a cluster

The YOPRO1 intensity profile was extracted and background subtracted. The estimated DNA content within each cluster was calculated as previously described.^35^ To estimate the number of Sox2 molecules in each cluster, we extracted the Cy5 intensity profile that colocalized with λDNA after subtraction of background signals. We then extracted the intensity profiles of Cy5-Sox2 non-specifically adsorbed to surface in the same field, which we deemed as monomers. The number of Sox2 molecules within each cluster on λDNA was estimated by dividing the cluster intensity by the monomer intensity.

#### Analysis of Pearson’s correlation coefficient

Time-averaged projection of the images in each channel was generated, and background was subtracted from each image. The regions of interest were segmented and extracted for further analysis. Pearson’s correlation coefficients in each condition were calculated using JaCoP plugin in FIJI.^36^ Costes’ randomized control, which describes the correlation between the randomly shuffled pixels of two compared images, was also calculated using the JaCoP plugin.

#### Condensation time analysis

Each immobilized λDNA molecule in a recorded field was individually monitored, and the time when a molecule condensed was defined as the transition at which the molecule completely lost slack/fluctuations. We subsequently ranked the condensation times and recorded the 75^th^ and 25^th^ percentile values (T_75%_and T_b_, respectively). The average condensation time (T_condense_) was calculated as T_75%_-T_25%_.

### Optical tweezers experiments

Single-molecule optical tweezers experiments were performed at room temperature on a LUMICKS C-trap combining confocal fluorescence microscopy and dual-trap optical tweezers, as previously described.^37^ In brief, we trapped two streptavidin-coated polystyrene beads (Spherotech) with a 1064-nm trapping laser and moved these beads to a channel containing biotinylated λDNA (LUMICKS). Single DNA tethers were selected based on the force-extension curve. The DNA tether was then moved into a channel containing 50 nM of Cy3-labeled Sox2 in T150 buffer. Cy3-Sox2 on DNA was visualized by confocal scanning with a 532-nm excitation laser. Correlative force and fluorescence measurements were made under different operation modes (force clamp mode, passive mode, or pulling mode)^38^ as specified in the figure legends.

### Electrophoretic mobility shift assay (EMSA)

DNA substrate was prepared via PCR and gel extraction of a 233-bp construct containing the Sox2 motif engineered into a 601 sequence, as previously described.^32^ 10 nM of DNA substrate was incubated with Sox2 and HMGB constructs in T150 buffer at room temperature for 30 min. The reaction mixture was loaded onto 5% non-denaturing polyacrylamide gel, which was run in 0.5× Tris-Borate-EDTA at 4°C at 100 V for 90 min, stained with SYBR Gold (Invitrogen), and visualized using a Typhoon FLA7000 gel imager (GE Healthcare).

### Statistical analysis

All error bars in this study represent 95% confidence intervals of the mean unless otherwise stated. *P* values were determined from unpaired two-tailed two-sample *t* tests (ns, not significant; * *P* <0.05, ** *P* <0.01, *** *P* <0.001).

## Figures

**Supplementary Fig. 1.**
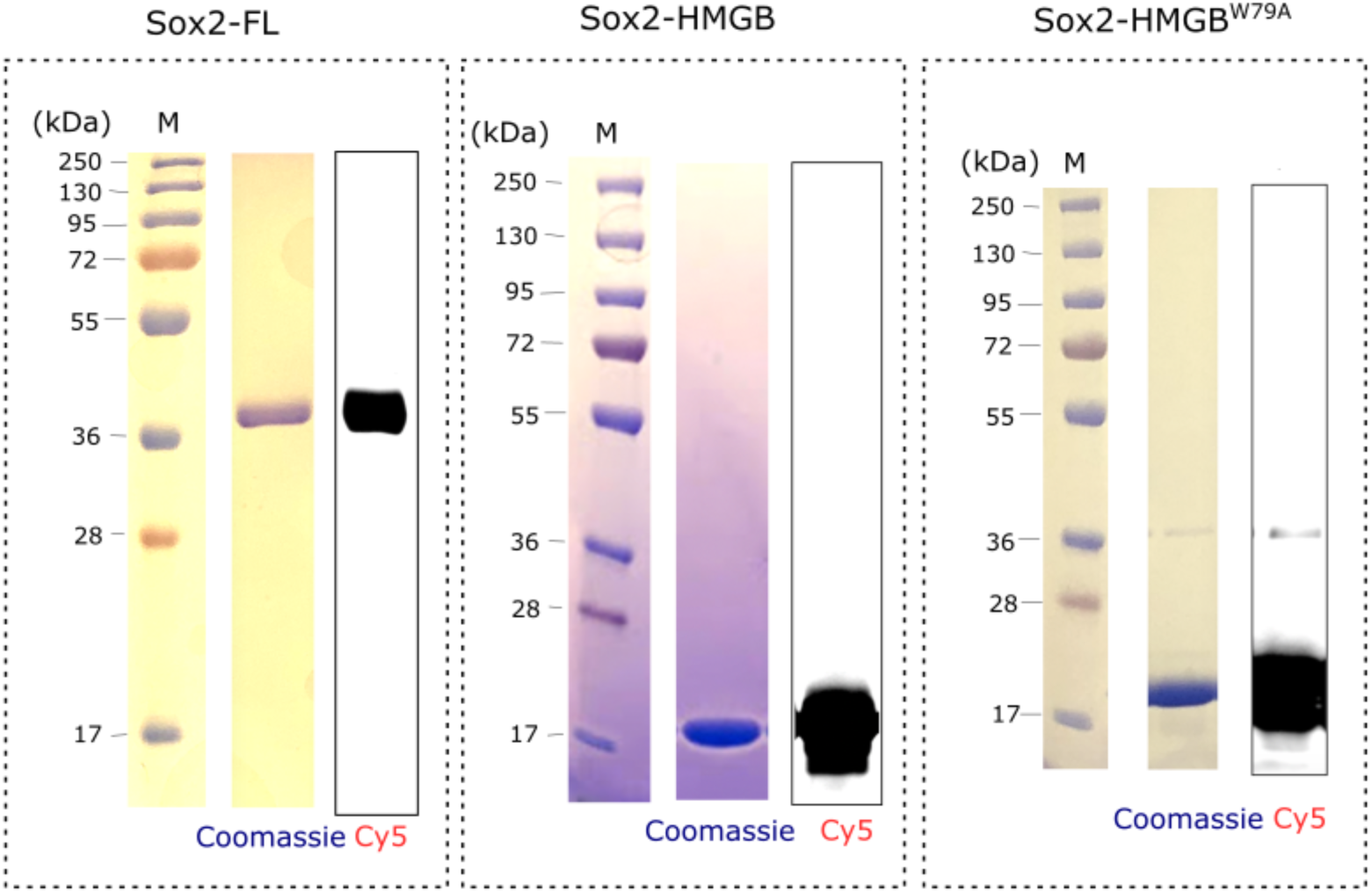
Purification and labeling of recombinant human Sox2. Coomassie stain and Cy5 fluorescence scan of Sox2-FL, Sox2-HMGB, and Sox2-HMGB^W79A^ proteins.

**Supplementary Fig. 2.**
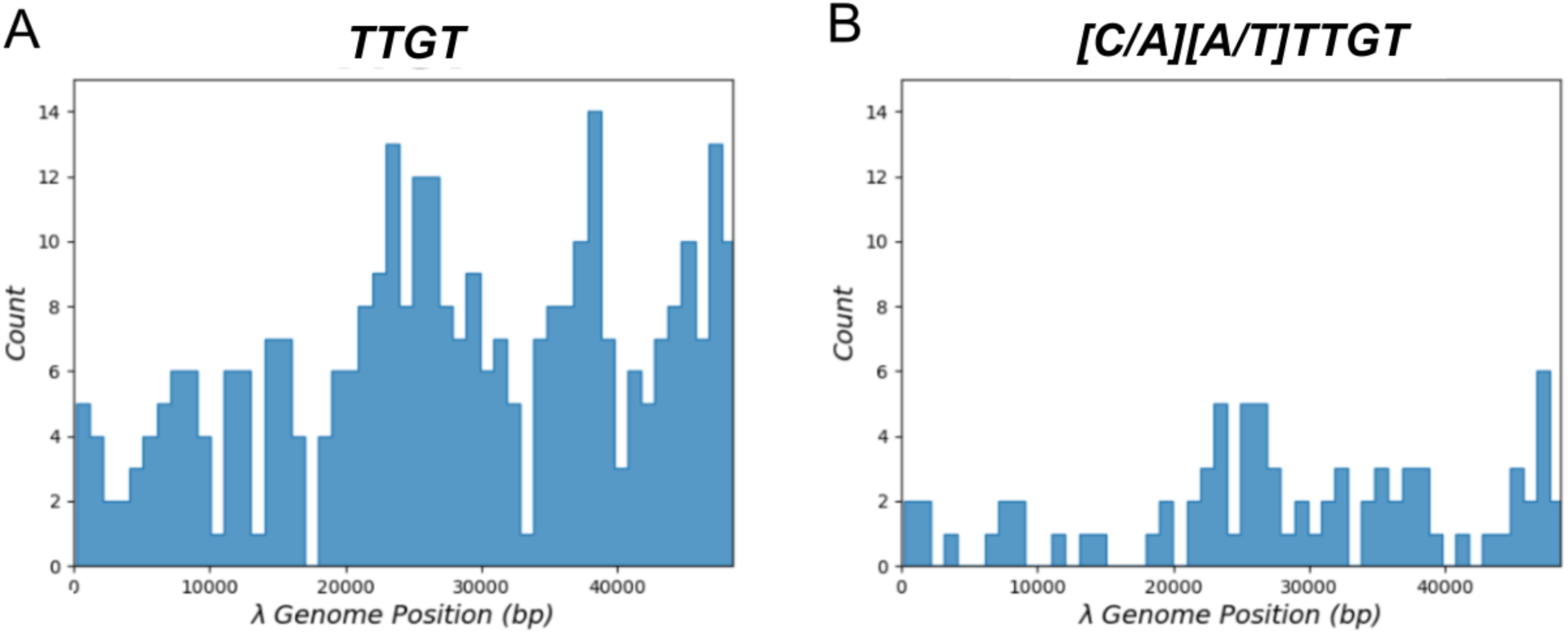
Distribution of Sox2 binding motifs on λDNA. A) Histogram displaying the occurrence of the canonical Sox2 motif TTGT along the λDNA genomic sequence.^39^ B) Histogram displaying the occurrence of the extended Sox2 motif [C/A][A/T]TTGT.^40^ Bin size in the histograms is 1 kb.

**Supplementary Fig. 3.**
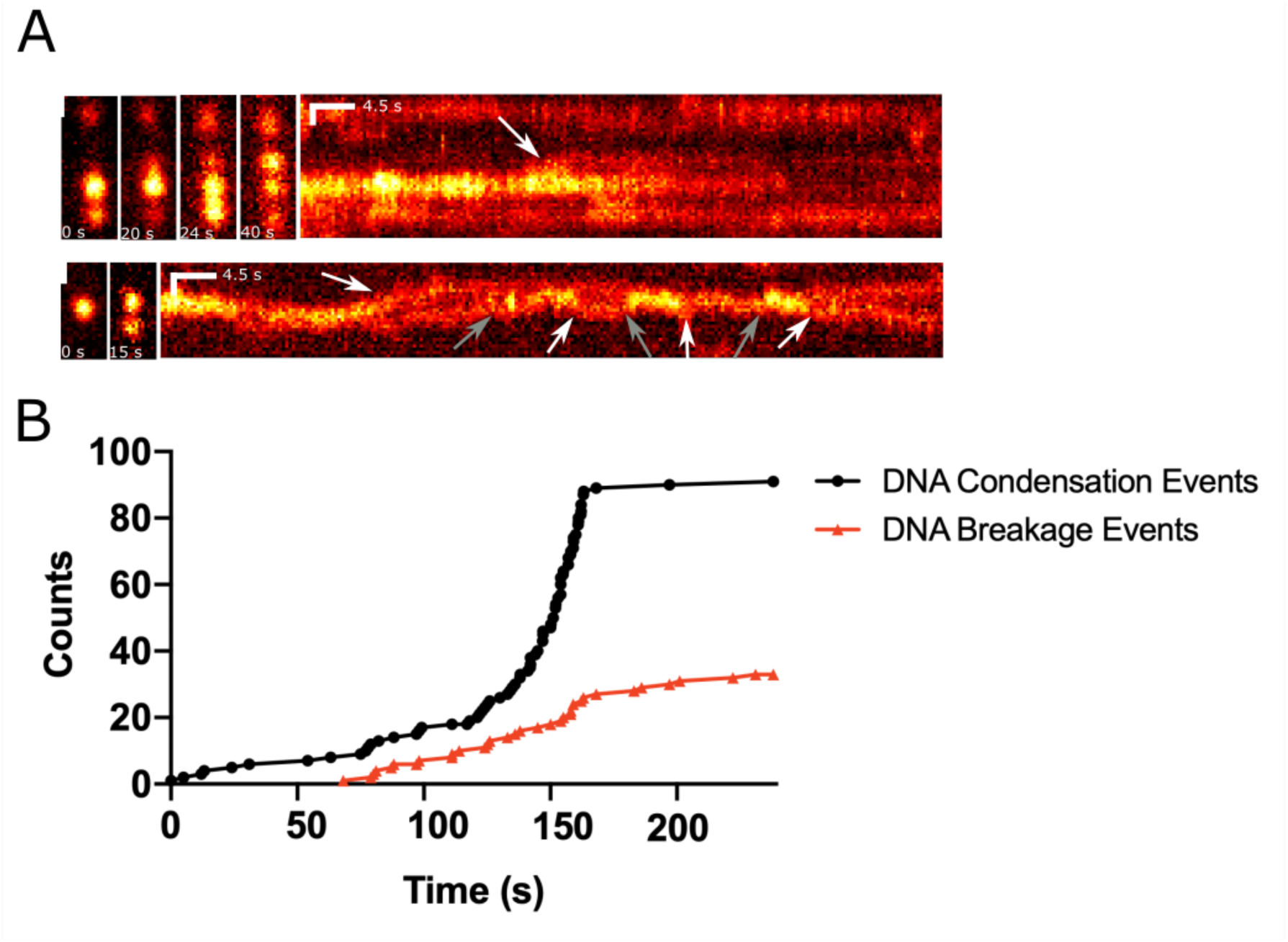
Additional characterizations of Sox2 condensates on DNA. A) Representative snapshots and kymographs of double-tethered λDNA molecules displaying fusion (gray arrows) and splitting events (white arrows) of Cy5-Sox2 foci. B) Cumulative incidence of Sox2-mediated DNA condensation and breakage events in a representative field of view.

**Supplementary Fig. 4.**
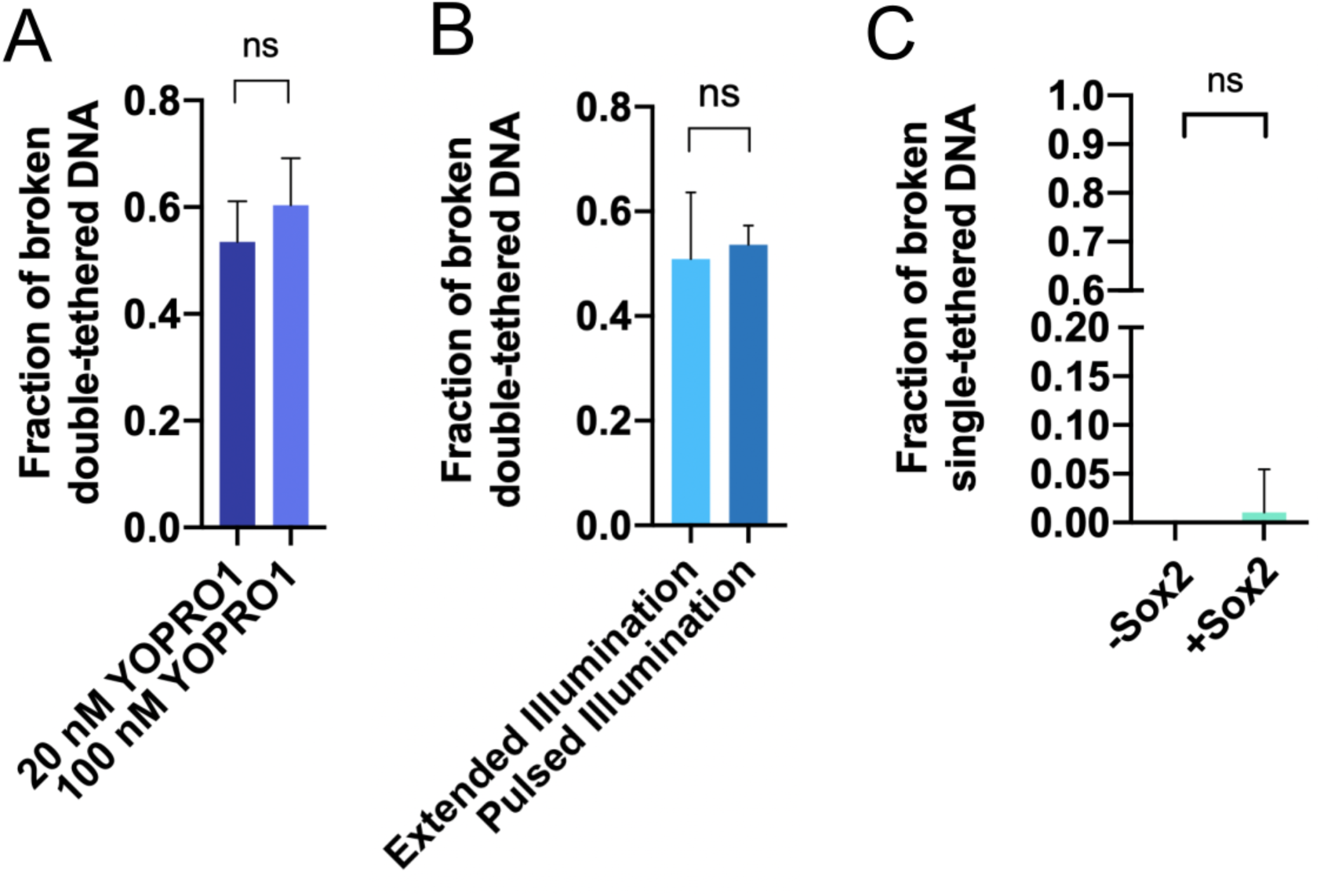
Control experiments supporting the model that Sox2 condensation is the main driver of DNA breakage. A) Fraction of double-tethered λDNA molecules that broke after 15 min of incubation with Sox2 in the presence of 20 nM (*n* = 157) or 100 nM (*n* = 106) YOPRO1. B) Fraction of double-tethered λDNA molecules in the presence of Sox2 that broke after 15 min under different laser illumination schemes. Under “Extended illumination”, 7.5 min of continuous 488-nm laser illumination was applied (*n* = 55). Under “Pulsed illumination”, a single 300-ms pulse of 488-nm laser at the same power was applied (*n* = 703). C) Fraction of single-tethered λDNA molecules that broke after 15 min of imaging in the absence of Sox2 (*n* = 165) or in the presence of Sox2 (*n* = 306). *n* represents the number of λDNA molecules analyzed.

**Supplementary Fig. 5.**
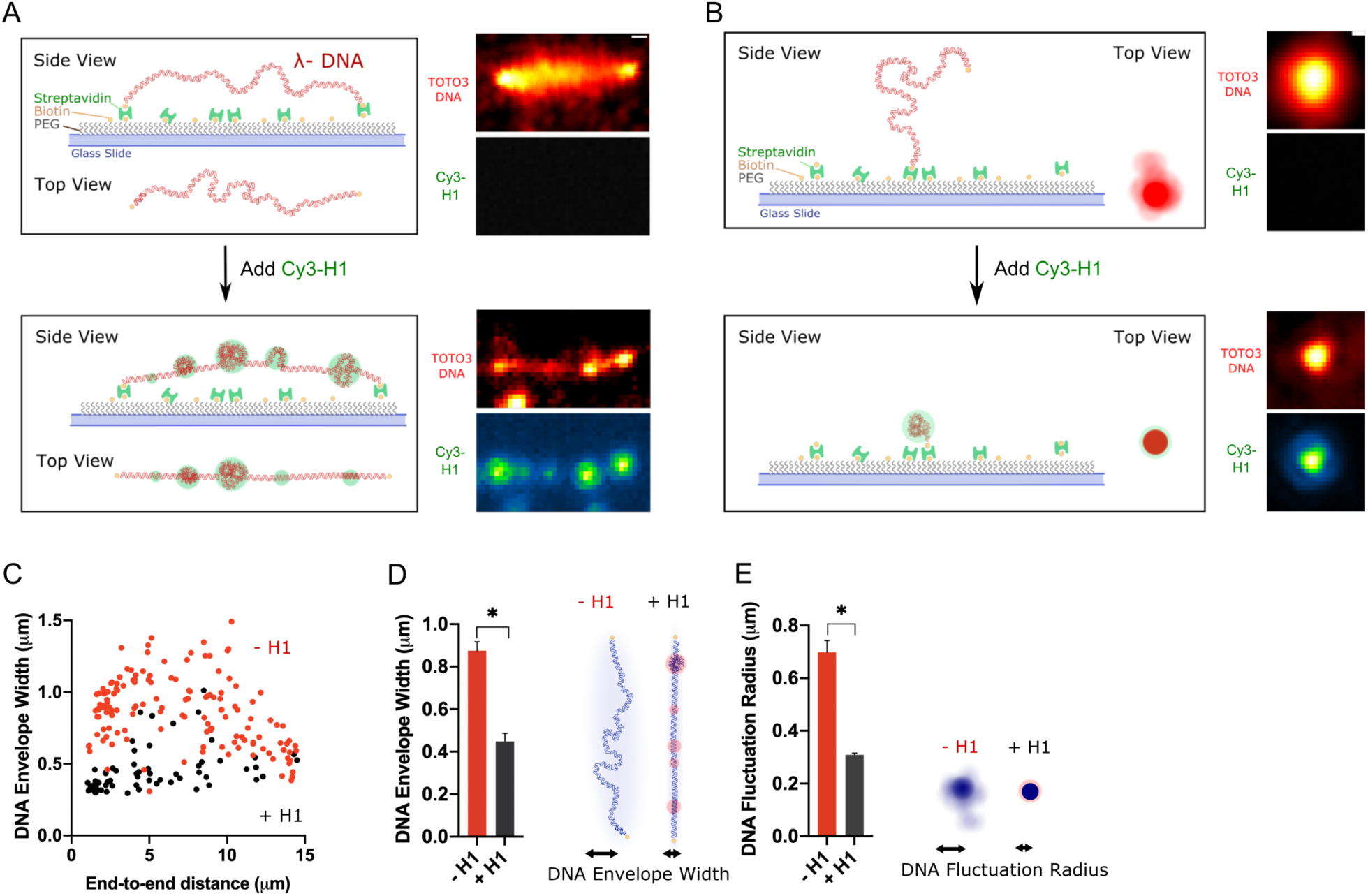
DNA condensation by linker histone H1. A) Schematic (left) and representative time-averaged projection (right) of a double-tethered λDNA stained with TOTO3 and incubated with 150 pM Cy3-labeled H1. Scale bar, 0.5 μm. B) Schematic (left) and representative time-averaged projection (right) of a single-tethered λDNA stained with TOTO3 and incubated with Cy3-H1. C) Double-tethered DNA envelope width as a function of end-to-end distance measured in the absence of H1 (red) and in the presence of H1 (black). D) Bar graph (left) and cartoon (right) showing a reduction in the average DNA envelope width of double-tethered λDNA upon H1-mediated DNA condensation (*n* = 63). E) Bar graph (left) and cartoon (right) showing a reduction in the average DNA fluctuation radius of single-tethered λDNA upon H1-mediated DNA condensation (*n* = 38).

**Supplementary Fig. 6.**
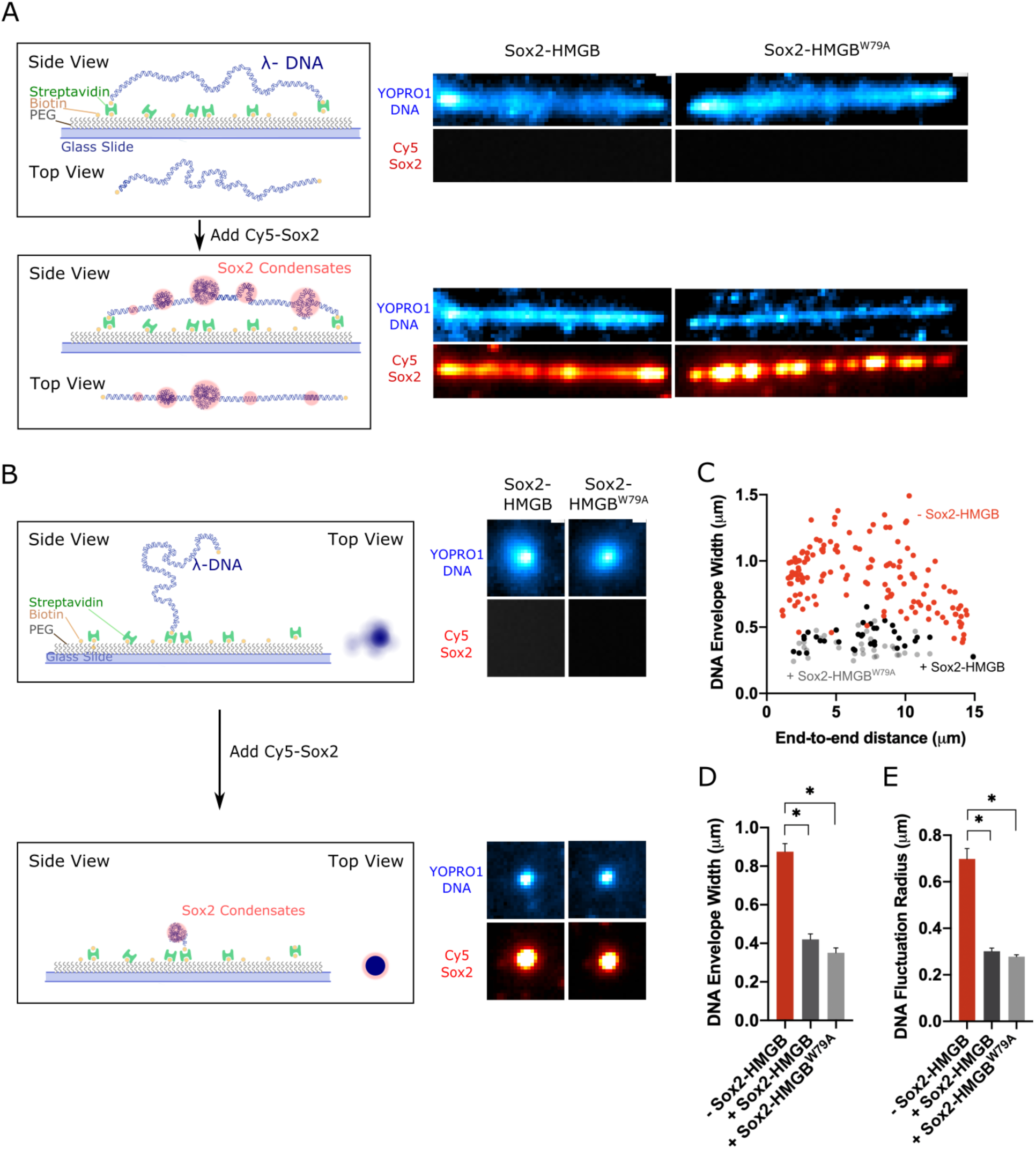
Additional results on DNA condensation mediated by Sox2-HMGB constructs. A) Schematic (left) and representative time-averaged projections (right) of double-tethered λDNA stained with YOPRO1 and incubated with 10 nM Cy5-labeled Sox2-HMGB or Sox2-HMGB^W79A^. Scale bar, 0.5 μm. B) Schematic (left) and representative time-averaged projections (right) of single-tethered λDNA stained with YOPRO1 and incubated with Cy5-labeled Sox2-HMGB or Sox2-HMGB^W79A^. C) Double-tethered DNA envelope width as a function of end-to-end distance measured in the absence of Sox2-HMGB (red) (*n*=149), in the presence of Sox2-HMGB (black) (*n* = 34), or Sox2-HMGB^W79A^ (gray) (*n* = 36). D) Bar graph showing a reduction in the average DNA envelope width of double-tethered λDNA molecules after incubation with Sox2-HMGB (*n* = 34) or Sox2-HMGB^W79A^ (*n* = 36). E) Bar graph showing a reduction in the average fluctuation radius of single-tethered λDNA molecules after incubation with Sox2-HMGB (*n* = 38) or Sox2-HMGB^W79A^ (*n* = 31).

**Supplementary Fig. 7.**
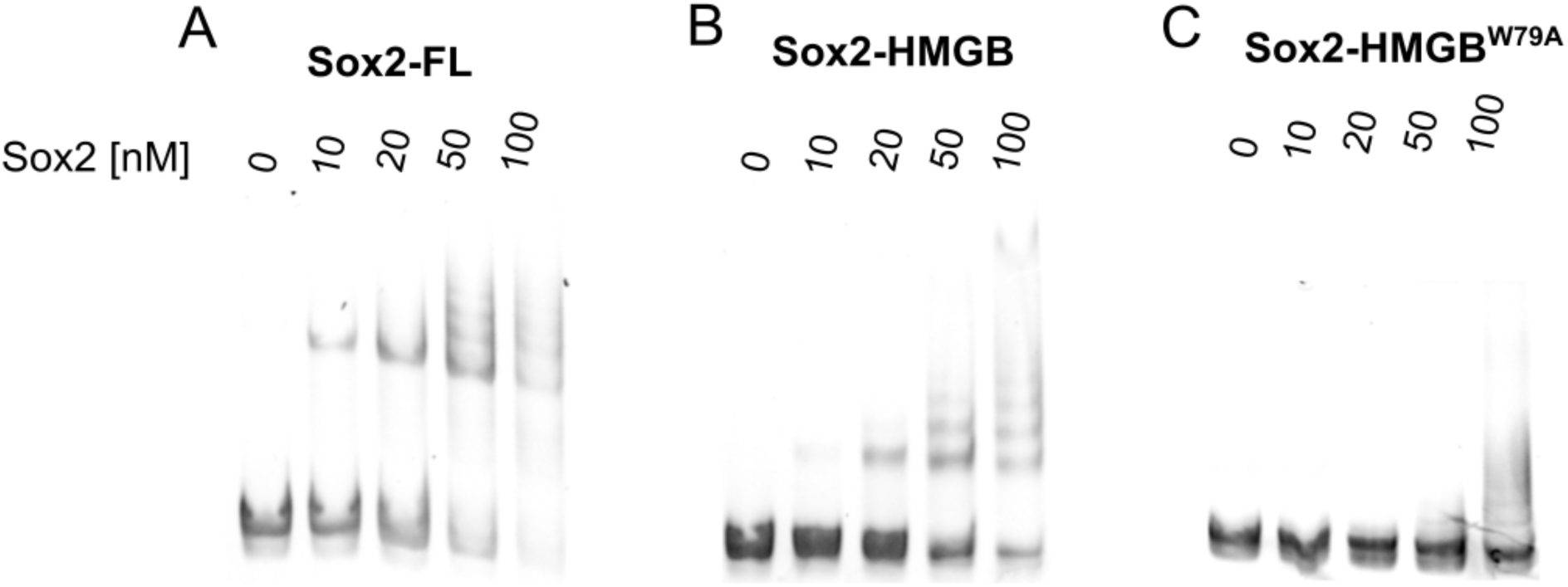
Electrophoretic mobility shift assay for the binding of different Sox2 constructs to DNA. A) SYBR-stained gel results for full-length Sox2 (Sox2-FL) binding to a 233-bp DNA that contains a Sox2 motif (CTTTGTT). B) Gel results for Sox2-HMGB binding to the same DNA. C) Gel results for Sox2-HMGB^W79A^ binding to the same DNA.

**Supplementary Fig. 8.**
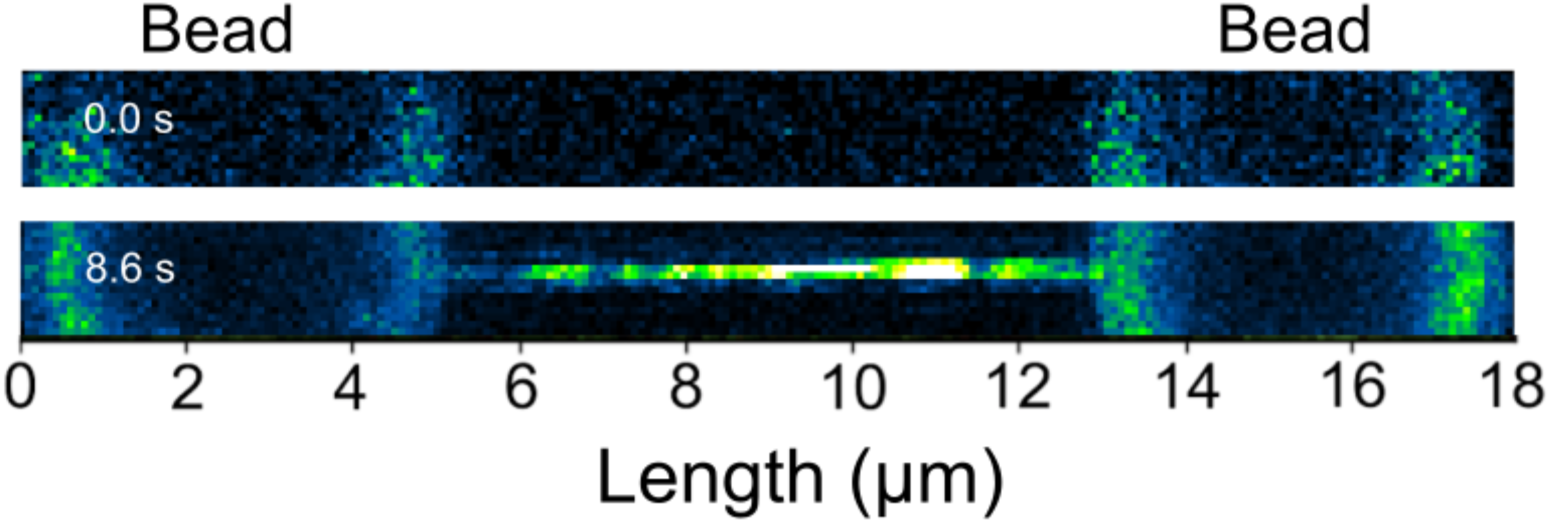
Visualization of Sox2 condensate formation on λDNA tethered between two optically trapped beads. Shown are 2D scans of a representative tether upon incubation with 50 nM Cy3-labeled Sox2. The trap positions were held fixed in this experiment. The initial force applied to the tether was close to zero. See Fig. 3B for force readings as a function of time.

**Supplementary Fig. 9.**
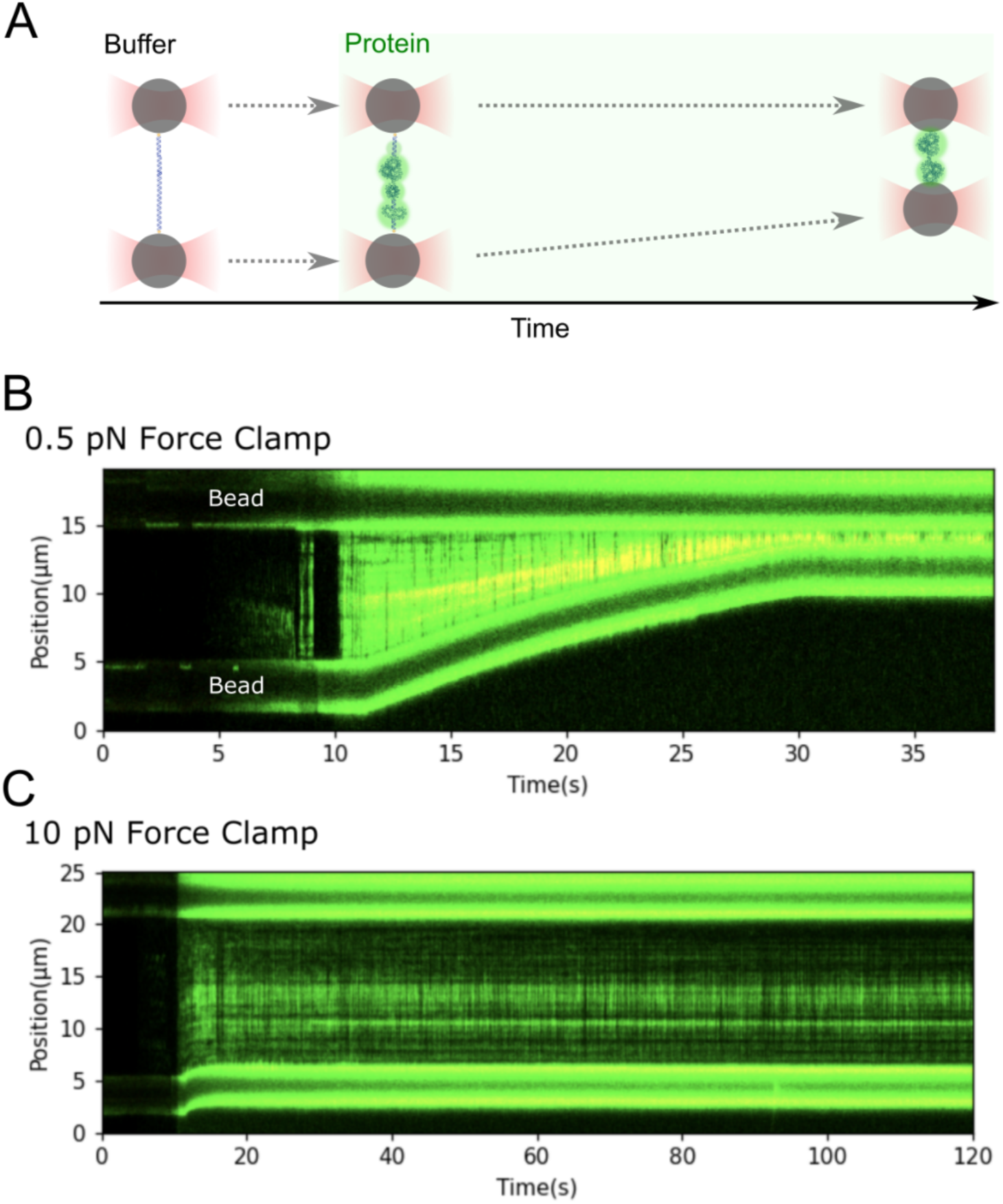
Evaluating the force-generating capability of Sox2-mediated DNA condensation with optical tweezers combined with fluorescence. A) Schematic of the optical tweezers assay. A λDNA molecule was tethered between two beads in a buffer channel and then moved to a Cy3-Sox2-containing channel. The force applied to the tether was kept constant via feedback (force clamp) such that DNA condensation would result in a shortening of the tether. B) A representative kymograph showing significant tether contraction and Sox2 condensate formation under a 0.5-pN force clamp. C) A representative kymograph showing suppressed tether contraction under a 10-pN force clamp.

**Supplementary Fig. 10.**
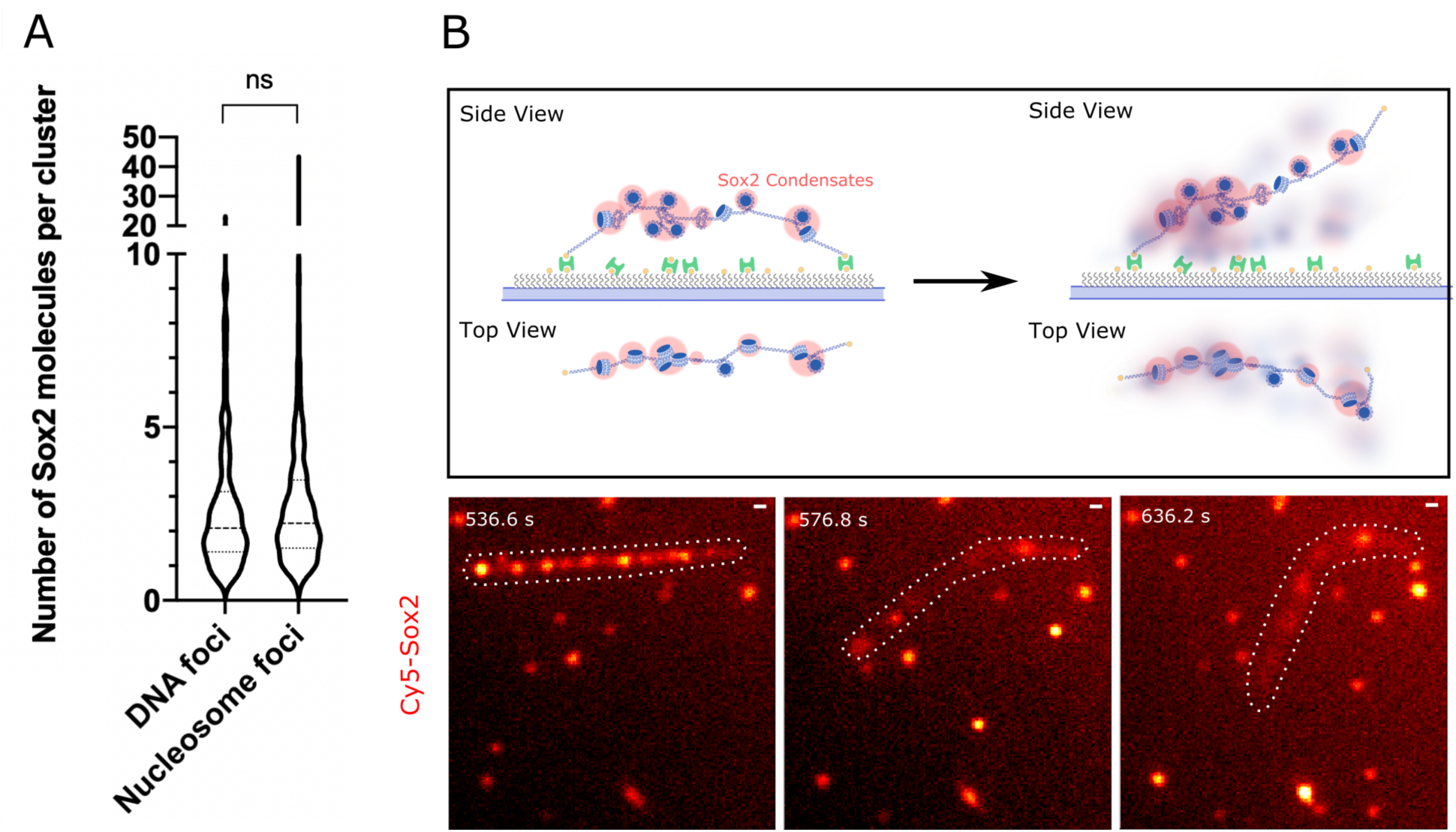
Sox2 binding and condensation on nucleosomal DNA. A) Violin plot showing the distribution of the number of Sox2 molecules within bare DNA foci (*n* = 381) versus within nucleosome foci (*n* = 616), where *n* represents the number of foci analyzed. B) Schematic (top) and representative snapshots (bottom) showing Sox2 condensates on nucleosomal DNA. Sox2-mediated condensation did not cause nucleosomal DNA to collapse in the same fashion as bare DNA. The nucleosomal DNA contour (white outlines) remained in an extended configuration and, when detaching from one anchored end, underwent rigid-body-like fluctuations. Scale bar, 0.5 μm. See also Supplementary Video 5.

## Supplementary Videos

**Supplementary Video 1**. An example showing Sox2-mediated suppression of single-tethered λDNA fluctuations. The DNA was stained with YOPRO1 (top) and Sox2 was labeled with Cy5 (bottom). The reduction of DNA fluctuation coincided with the increase of Sox2 signal (~ 17 sec). Scale bar: 0.5 μm.

**Supplementary Video 2**. An example showing that the formation of Sox2 condensates on double-tethered λDNA causes a reduction in transverse tether fluctuations and eventually DNA breakage. The DNA was stained with YOPRO1 (top) and Sox2 was labeled with Cy5 (bottom).

Scale bar: 0.5 μm.

**Supplementary Video 3**. An example showing that Sox2 condensate formation mediates the joining of nearby DNA strands. The DNA was stained with YOPRO1 (top) and Sox2 was labeled with Cy5 (bottom). Scale bar: 0.5 μm.

**Supplementary Video 4**. An example demonstrating the mechanical effects of Sox2-mediated DNA condensation in *cis* and *trans*. Multiple DNA breaking and joining events were observed. The DNA was stained with YOPRO1 for fluorescence visualization. See Fig. 2D for snapshots and cartoon illustrations.

**Supplementary Video 5**. An example showing rigid-body fluctuations of nucleosomal DNA (visualized by the fluorescence signal from bound Cy5-Sox2) after one end detached from the surface. See Supplementary Fig. 9B for snapshots and cartoon illustrations. Scale bar: 0.5 μm.

**Supplementary Video 6**. An example showing fluctuations of the same single-tethered nucleosomal λDNA that persist in the presence of Sox2 (left: without Sox2; right: with Sox2). The nucleosomal DNA was visualized by the fluorescence signal from Cy3-H2B (top) and Sox2 was labeled with Cy5 (bottom). Scale bar: 0.5 μm.

All videos are played at 10× speed.

## References

1. Ptashne, M. & Gann, A. Genes & Signals. (Cold Spring Harbor Laboratory Press, 2002).

2. Boija, A. et al. Transcription Factors Activate Genes through the Phase-Separation Capacity of Their Activation Domains. Cell 175, 1842–1855 (2018).

3. Chong, S. et al. Imaging dynamic and selective low-complexity domain interactions that control gene transcription. Science 361, eaar2555 (2018).

4. Shin, Y. et al. Liquid Nuclear Condensates Mechanically Sense and Restructure the Genome. Cell 175, 1481–1491 (2018).

5. Larson, A. G. et al. Liquid droplet formation by HP1α suggests a role for phase separation in heterochromatin. Nature 547, 236–240 (2017).

6. Hnisz, D., Shrinivas, K., Young, R. A., Chakraborty, A. K. & Sharp, P. A. A Phase Separation Model for Transcriptional Control. Cell 169, 13–23 (2017).

7. Cramer, P. Organization and regulation of gene transcription. Nature 573, 45–54 (2019).

8. Xiao, B., Freedman, B. S., Miller, K. E., Heald, R. & Marko, J. F. Histone H1 compacts DNA under force and during chromatin assembly. Mol. Biol. Cell 23, 4864–4871 (2012).

9. Bell, N. A. W. et al. Single-molecule measurements reveal that PARP1 condenses DNA by loop stabilization. Sci. Adv. 7, eabf3641 (2021).

10. Leicher, R. et al. Single-stranded nucleic acid sensing and coacervation by linker histone H1. bioRxiv (2021).

11. Renger, R. et al. Co-condensation of proteins with single- and double-stranded DNA. bioRxiv (2021).

12. Feric, M. Droplets take DNA by force. Nat. Phys. 17, 981–982 (2021).

13. Quail, T. et al. Force generation by protein – DNA co-condensation. Nat. Phys. 17, 1007– 1012 (2021).

14. Luger, K., Dechassa, M. L. & Tremethick, D. J. New insights into nucleosome and chromatin structure: an ordered state or a disordered affair? Nat. Rev. Mol. Cell Biol. 13, 436–447 (2012).

15. Hock, R., Furusawa, T., Ueda, T. & Bustin, M. HMG chromosomal proteins in development and disease. Trends Cell Biol. 17, 72–79 (2007).

16. Xiao, B., Freedman, B. S., Miller, K. E., Heald, R. & Marko, J. F. Histone H1 compacts DNA under force and during chromatin assembly. Mol. Biol. Cell 23, 4864–4871 (2012).

17. Krainer, G. et al. Reentrant liquid condensate phase of proteins is stabilized by hydrophobic and non-ionic interactions. Nat. Commun. 12, 1085 (2021).

18. Xue, B., Oldfield, C. J., Van, Y., Keith, A. & Uversky, V. N. Protein intrinsic disorder and induced pluripotent stem cells. Mol. Biosyst. 8, 134–150 (2012).

19. Holmes, Z. E. et al. The Sox2 transcription factor binds RNA. Nat. Commun. 11, 1805 (2020).

20. Soufi, A. et al. Pioneer transcription factors target partial DNA motifs on nucleosomes to initiate reprogramming. Cell 161, 555–568 (2015).

21. Banigan, E. J. & Mirny, L. A. Loop extrusion: theory meets single-molecule experiments. Curr. Opin. Cell Biol. 64, 124–138 (2020).

22. Morin, A. J. A. et al. Surface condensation of a pioneer transcription factor on DNA. bioRxiv (2020).

23. Keenen, M. M. et al. HP1 proteins compact DNA into mechanically and positionally stable phase separated domains. Elife 10, e64563 (2021).

24. Bustamante, C., Chemla, Y. R., Forde, N. R. & Izhaky, D. Mechanical Processes in Biochemistry. Annu. Rev. Biochem. 73, 705–748 (2004).

25. Bico, J., Reyssat, É. & Roman, B. Elastocapillarity: When Surface Tension Deforms Elastic Solids. Annu. Rev. Fluid Mech. 50, 629–659 (2018).

26. Sharma, R. et al. Liquid condensation of reprogramming factor KLF4 with DNA provides a mechanism for chromatin organization. Nat. Commun. 12, 5579 (2021).

27. Courchaine, E. M. et al. DMA-tudor interaction modules control the specificity of in vivo condensates. Cell 184, 3612–3625 (2021).

28. Bystricky, K., Heun, P., Gehlen, L., Langowski, J. & Gasser, S. M. Long-range compaction and flexibility of interphase chromatin in budding yeast analyzed by high-resolution imaging techniques. Proc. Natl. Acad. Sci. U. S. A. 101, 16495–16500 (2004).

29. Le, T. T. et al. Synergistic Coordination of Chromatin Torsional Mechanics and Topoisomerase Activity. Cell 179, 619–631 (2019).

30. Zaret, K. S. & Carroll, J. S. Pioneer transcription factors: Establishing competence for gene expression. Genes Dev. 25, 2227–2241 (2011).

31. Miroshnikova, Y. A., Nava, M. M. & Wickstro, S. A. Emerging roles of mechanical forces in chromatin regulation. J. Cell Sci. 130, 2243–2250 (2017).

32. Li, S., Zheng, E. B., Zhao, L. & Liu, S. Nonreciprocal and Conditional Cooperativity Directs the Pioneer Activity of Pluripotency Transcription Factors. Cell Rep. 28, 2689–2703 (2019).

33. Osunsade, A. et al. A Robust Method for the Purification and Characterization of Recombinant Human Histone H1 Variants. Biochemistry 58, 171–176 (2018).

34. Crickard, J. B., Moevus, C. J., Kwon, Y., Sung, P. & Greene, E. C. Rad54 Drives ATP Hydrolysis-Dependent DNA Sequence Alignment during Homologous Recombination. Cell 181, 1380–1394 (2020).

35. Ganji, M. et al. Real-time imaging of DNA loop extrusion by condensin. Science 360, 102– 105 (2018).

36. Bolte, S. & Cordelieres, F. P. A guided tour into subcellular colocalization analysis in light microscopy. J. Microsc. 224, 213–232 (2006).

37. Wasserman, M. R. et al. Replication Fork Activation Is Enabled by a Single-Stranded DNA Gate in CMG Helicase. Cell 178, 600–611 (2019).

38. Bustamante, C. J., Chemla, Y. R., Liu, S. & Wang, M. D. Optical tweezers in single-molecule biophysics. Nat. Rev. Methods Prim. 1, 25 (2021).

39. Schaefer, T. & Lengerke, C. SOX2 protein biochemistry in stemness, reprogramming, and cancer : the PI3K / AKT / SOX2 axis and beyond. Oncogene 39, 278–292 (2020).

40. Hou, L., Srivastava, Y. & Jauch, R. Molecular basis for the genome engagement by Sox proteins. Semin. Cell Dev. Biol. 63, 2–12 (2017).

